# Do Cells use Passwords in Cell-State Transitions? Is Cell Signaling sometimes Encrypted?

**DOI:** 10.1101/432120

**Authors:** Alex Root

## Abstract

Organisms must maintain proper regulation including defense and healing. Life-threatening problems may be caused by pathogens or by a multicellular organism’s own cells through cancer or auto-immune disorders. Life evolved solutions to these problems that can be conceptualized through the lens of information security, which is a well-developed field in computer science. Here I argue that taking an information security view of cells is not merely semantics, but useful to explain features of signaling, regulation, and defense. An information security perspective also offers a conduit for cross-fertilization of advanced ideas from computer science, and the potential for biology to inform computer science. First, I consider whether cells use passwords, i.e., initiation sequences that are required for subsequent signals to have effects, by analyzing the concept of pioneer transcription factors in chromatin regulation and cellular reprogramming. Second, I consider whether cells may encrypt signal transduction cascades. Encryption could benefit cells by making it more difficult for pathogens or oncogenes to hijack cell networks. By using numerous molecules cells may gain a security advantage in particular against viruses, whose genome sizes are typically under selection pressure. I provide a simple conceptual argument for how cells may peform encryption through post-translational modifications, complex formation, and chromatin accessibility. I invoke information theory to provide a criterion of an entropy spike to assess whether a signaling cascade has encryption-like features. I discuss how the frequently invoked concept of context-dependency may over-simplify more advanced features of cell signaling networks, such as encryption. Therefore, by considering that biochemical networks may be even more complex than commonly realized we may be better able to understand defenses against pathogens and pathologies.

## Main

Biochemical networks transmit, receive, and process information resulting in decision making concerning growth, defense, differentiation, migration, apoptosis, metabolism, and other processes^1,2^. Groundbreaking studies over the last several decades have elucidated properties of signaling and regulatory networks, including scale-freeness, robustness, fragility, noise-filtering, bistability, controllability, ultrasensitivity, signal dissipation, amplification, memory, modularity, feedfoward and other motifs, which are reviewed by Krakauer and colleagues^2^, Uda and Kuroda^3^, Mousavian and colleagues^4,5^, Walterman and Klipp^6^, Azeloglu and Iyengar^1^, and Antebi and colleagues^7^. Biochemical networks can become dysfunctional through genetic mutation, chemical injury, infection, or other processes, that might achieve varying degrees of control over the network^8^. Here, I consider these processes through the lens of information security, which as far as I can determine is not common. This is in contrast to telecommunications, where cybersecurity is of paramount importance^9^. In an elegant and trenchant examination of theoretical biology, Krakauer and colleagues argue “before we can look for patterns, we often need to know what kinds of patterns to look for, which requires some fragments of theory to begin with^10^.”

By explicitly incorporating information security concepts into thinking about biological systems, several outcomes are possible in general: (1) distinctions without differences, that is, we are merely rephrasing familiar concepts of immunity and regulation in terms of information security, which adds no value; (2) cross-disciplinary fertilization occurs as information security concepts are imported into biological theory; (3) new information security knowledge arises from examination of biological systems. Recent studies on network controllability provide one framework for examining security in biochemical networks^11–15^. Here, a different perspective is taken to analyze whether cells may use passwords and encrypt their signaling cascades.

## Immune systems and biological security

The evolution of immune systems and self-defense against injury and mutation are major innovations in the history of life on earth^16–18^. By total volume, life on earth has its largest habitat in the deep ocean with an abundance of bacteriophages, seemingly saturating the environment^19^. Indeed, parasitism has been estimated to have evolved independently several hundred times. Single-celled and multi-cellular organisms evolved a wide-variety of defense systems, often dichotomized into innate and adaptive systems^20^. These systems can be conceptualized more generally to include protective mechanisms against both external and internal damage. The connection between external and internal injury is seen in the study of viruses, which led to insights in cancer biology and the discovery of oncogenes^17^. Organisms developed the ability to recognize self from non-self and destroy xenobiotic material. However, not all foreign genetic material is completely destroyed, because it can increase fitness, e.g., antibiotic resistance plasmids^20,21^. On the intracellular level, bacterial defense mechanisms include blocking receptor binding (surface modification), genome injection (superinfection exclusion), viral replication (restriction modification, CRISPR-Cas, and prokaryotic Argonaute), and abortive infection (programmed cell death)^21^. Similar mechanisms exist in eukaryotic cells, including, RIG-like receptor proteins that recognize RNA^16^, xenophagy^22^, advanced intracellular nucleic acid recognition systems and other cell-autonomous mechanisms^23^. In plants, sophisticated DICERs defend against retroviruses^24^. Similarly, pathogens use a variety of mechanisms to co-opt, hijack, and counteract host defenses^25–28^. Mutations leading to oncogenes reprogram signaling networks^29^. All of these attacks and counter-attacks involve changes in signaling and regulatory networks, and therefore, changes in information. Intriguingly, the ciliates *Oxytricha* and *Stylonychia* have scrambled or encrypted genomes that undergo massive re-arrangement during development into functional genes^30^. Presumably, this evolved as a defense mechanism against pathogens and provides examples that cells encrypt information.

## Information security in computer science

Information security has been important for millennia, with the Caesar substitution cipher being a known early example^31^. (The cipher works by shifting each letter of the English alphabet by 3, i.e., A->C, B->D,…,X->A^31^). Within several decades of the development of computers, security against computer viruses became an important concern. Computer viruses achieved notoriety in 1987 when the Brain, Lehigh, and April Fool viruses came to worldwide attention^32^. Hackers achieved infamy and also contributed to the advancement of information technology^33^. Information security depends on the use of passwords for system access and encryption^34^. Development of secure encryption systems, e.g., the RSA asymmetric public key cryptography, was an essential innovation in the history of the internet^34^ and must evolve to meet new threats^9^. Steganography is an altogether different approach that conceals the existence of information, e.g., writing with invisible ink, and appears to have had played less importance in the history of information technology than cryptography^34^. Attacks on encrypted systems can involve interception, modification, fabrication, or interruption of information^34^. There has been considerable work in adapting biomolecules for use in information security in human telecommunications using biosteganography^35^ where information is invisible and molecular cryptography, where synthetic biology is used to re-engineer molecules to decode and encode information^36^. Despite obvious parallels to computer science, less explicit attention appears to have been paid to theoretical descriptions of cells in terms of information security.

## Information systems in cells

Organisms have a variety of sophisticated information systems. They encode information through the nucleic acid sequences, which utilize complementary base pairing of two strands to provide built-in error correction, which is a type of backup and repair system. At the proteome level, cells can modify amino acids with a few hundred post-translational modifications in various combinations, that give rise to numerous proteoforms^37^, which form components of signaling and regulatory networks. Somatic recombination in immunoglobulins and T-cell receptors can vastly increase protein variants in certain cell types^38^. Interactions of these macromolecules form networks that store and transmit information^6^. There is a context specificity to many signaling pathways, including TGF-beta and AKT, which means that cells respond differently to pathway activation depending on the cell type^39,40^. Many intracellular signaling pathways do not match one receptor to a single ligand, but instead use multiple receptors and ligands that interact combinatorially^41^, or use combinations of numerous nuclear-receptor cofactors to regulate activity^42^. Therefore, genetic, epigenetic, transcriptomic and proteomic variation gives rise to a large repertoire of interacting components. These mechanisms are present in complex multicellular organisms, where advanced regulation is needed to control differentiation^43^ and also in bacteria for complex behaviors, such as quorum sensing^2^.

Cancer often involves rewiring cellular networks by oncogenes and therefore, in some sense, these represent alterations in information transmission and compromised security^29,44^. Cancer cells can transition because of the effects of microRNAs and oncogenes to different states with distinct metabolism^45^. Similarly, viruses can substantially rewire signaling and regulatory networks to hijack cellular machinery for viral benefit^46^. In the early days of cancer research, similarities between the two systems caused the scientific community to think that viruses cause cancer, and studies into viral biology provided insights into cancer^17,47^. Both pathogenic and pathological processes can involve hijacking cellular networks.

In multicellular organisms, combinations of histone modifications give rise to varying chromosomal accessibility and epigenetic states, which are read, written, and erased^48,49^. This epigenetic regulation is capable of encoding memory at the single-cell level^50^. Redundancy and correlation among epigenetic marks, transcription factors, and co-regulators provides a way to specify cell states, off which there are more than 250 distinct cell types in adult humans^51^. For example, the identity of a ligand can be encoded as pulsatile (DLL1-Notch1) or sustained (DLL4-Notch1) to induce opposite cell fates. In the adult human body, several hundred distinct cell types exist in cell states, some of which can be transitioned from one state to the next using combinations of growth factors and transcription factors^52–54^. The language used to describe these cellular properties (code, encode, read, write, memory, erase, rewire) suggests that they may be information systems, but requires careful thought to address *how* it is that they are information systems.

## Do cells use passwords before transitioning between cell states?

Password authorization systems allow access based upon entry of a somewhat random code and achieve security by creating a large search space. Passwords are also an initiation sequence of signals without which a system will not respond to subsequent signals. In multicellular organisms, cells develop into types that can be conceptualized as stable states. In order to change from one state to another, that is, to differentiate or de-differentitate, the epigenome of cells changes^55^.

Organization of chromatin into highly compact, inaccessible regions, and open, accessible regions is a form of information security because some genes are “locked” and therefore, cannot be transcribed. Chromatin is frequently characterized as being in “open” and “closed” states that must be unlocked. This appears to be a potential case where cells use passwords because the right combination of growth factors and transcription factors are necessary for reprogramming to occur. The initial sequence is mediated by so-called pioneer transcription factors. There are multiple algorithms to predict combinations of transcription factors to reprogram human cells from one type to another with the number of successful conversions being relatively low^52^. The systems work by engineering overexpression of transcription factors, rather than as it happens normally in development through extracellular signaling molecules that signal to transcription factors to achieve the rewiring. To my knowledge, no one has attempted to predict upstream combinations of signals, e.g., growth factors, hormones, adhesion contacts, etc. that would trigger the right combinations of transcription factors. Sampattavanich and colleagues demonstrated that FOXO3 dynamics can code for different growth factors and their concentrations, which are under combinatorial control of ERK and AKT pathways^56^. Typically, engineering the over-expression of transcription factors does not introduce dynamics, that is, the over-expression simply turns on to high concentration.

If cell reprogramming requires an initiation sequence of transcription factors to unlock the epigenetic code then conceptualizing this as a password is appropriate. A password conceptualization is distinct from simply requiring a series of events because of its defensive purpose and strategy of employing a seemingly random draw from a large search space. If a set of transcription factors are active during the entire reprogramming process, then they are not performing an initiation sequence and therefore, not entering a password. Similarly, if only one member of the combination can partially reprogram cells then it would seem inappropriate to conceptualize the mechanism as a password. This conceptualization might predict that password-length, i.e., the complexity of the reprogramming initiation is directly proportional the fitness cost posed to the organism from the conversion. For example, because stem cells have greater replicative potential, they might pose greater risk to develop into cancer and consequently, require a more complex password for reprogramming. It is unclear to me whether there are sufficient measurements to resolve whether there is a distinct initiation sequence during cell reprogramming.

## Do cells encrypt information in signaling cascades?

Cell signaling can be hijacked by pathogens or oncogenes; for example, the chrysanthemum virus B p12 protein acts as a transcription factor to control cell size and proliferation for its benefit^57^. A cell signaling system might be able to use an encryption-like system to block this sort of attack. The way that it might work is to require different combinations of signals in different cell types to control cell size and proliferation. What encryption systems do is to increase the entropy of the signal by having machinery to encrypt and decrypt, e.g., rotors on an enigma machine. As a general rule, there is evolutionary pressure on viruses to maintain relatively small genomes, and less constraint on the sizes of non-viral genomes. This gives hosts a competitive advantage over viruses to create more complex signaling systems by adding components, e.g., encrypters/decrypters, in the form of additional genes. It is also protection against deleterious mutations because these are less likely to occur in multiple genes. The cost would seem to be maintaining a larger genome.

One way encryption could work in cell signaling is through a system consisting of an input signal, an encoder, an encrypter, a noisy signal transduction cascade, a decrypter, a decoder, and an output signal. Hypothetically, an encrypter could consist of multiple post-translational modifications on cell surface receptors and a decrypter could consist of transcription factor complexes interacting with post-translationally modified histones, see figure 1A. For example, suppose two different cell types A and B receive the same binary input signal, which they decode identically to the same output signal, such as a cascade of gene expression to differentiate into a another cell type. Suppose the phosphorylation patterns differ on the receptors, the transcription factor complexes differ, and the chromatin state on the target genes differ, see figure 1. The conventional explanation for these different signaling cascades is context-dependence, i.e., different signals in different cell types. Context-dependence is itself a useful form of information security because a different attack may be required for each context, so that only a subset of an organisms’ tissues may be attacked at any one time. What is needed to distinguish context-dependence from encryption is a consideration of what encryption does quantitatively, that is, encryption causes an increase in entropy of the transmitted signal, see figure 1B. Figure 1A depicts this as an increase in possible post-translational modification states going from the encoder (receptor binding on/off) to encrypter (receptor dimerization partner with 3 on/off phosphorylation sites (8 states)), where the inactive pathway randomly transmits the various states. An attacker is therefore less likely to mimic the activation signal during the encrypted chain, especially if the chain has dynamic behavior, such as periodic oscillations, see figure 1C. However, vulnerabilities are still present before encryption and after decryption. This suggests that a more secure biochemical system would combine signal reception, encoding, and encryption into an integrated ligand-receptor-scaffold complex. Or it suggests that there may be something about the biochemistry of the input and output signals that is harder for a pathogen or pathology to mimic, e.g., an extracellular ligand when typically viruses infect intracellularly.

**Figure 1.**
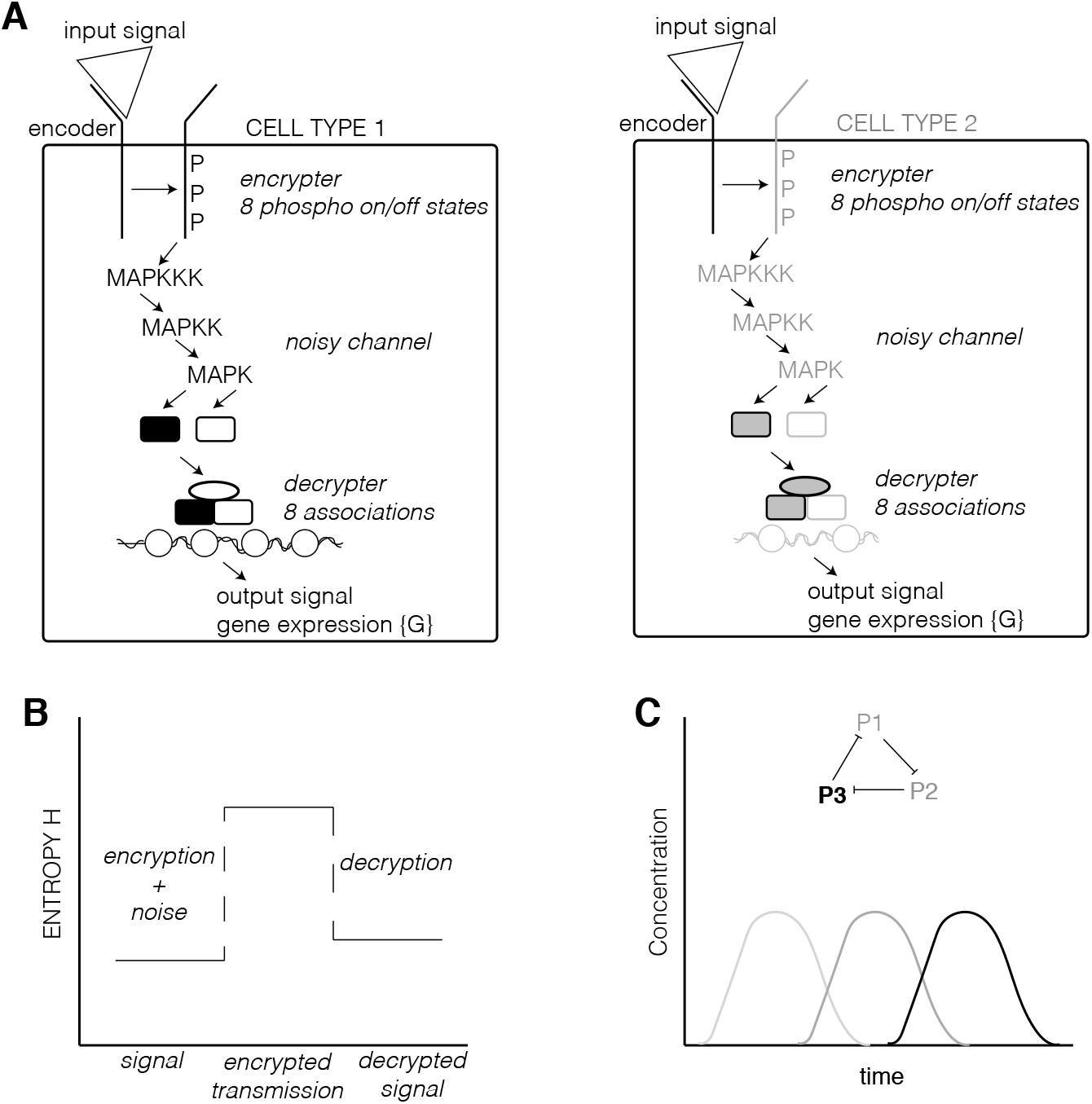
How encryption might work in cell signaling. **A** Two different cell types encode and encrypt the same signal binary signal of ligand binding using different receptor dimerization pairs with 3 phosphorylation sites, allowing for 8 on/off states, only a subset of which encode the signal and the remainder are transmitted randomly when the receptor is inactive. The encrypted signal is relayed through a noisy channel where a complex of transcription factors and cofactors can also form 8 different association states, only a subset of which can interact with the chromatin state of the cell to produce an output response of expresed genes. **B** If an input signal is encrypted then its transmission will have a higher entropy. The decrypted output signal will have an entropy slightly higher than the input signal due to noise. The presence of error correction may make noise minimal. **C** The repressilator circuit can produce periodic oscillations in a biochemical system that in principle could function like a substitution cipher, i.e., A->A, A->B, A->C, A->A, etc.

## Conclusions

Evolutionary potential is vast and a complex interplay among environmental change, ecosystems, speciation, niche diversification, extinctions, and innovation have shaped life on earth^58,59^. Considering how rapidly hacking, passwords, and encryption evolved in computers, it is natural to ask whether they are used in nature by cells. This conceptual exploration suggests that cells may use passwords to lock-in cell state, which must be unlocked through the right combination of transcription factors. Open questions include if cells use passwords to initiate cell signaling cascades, programmed cell death, neuron-to-neuron transmissions, or other areas. Also, is it the case that password-length, i.e., the complexity of an initiation sequence is directly proportional the fitness cost? When there is selection pressure due to co-evolution of pathogens are there more complex initiation sequences, i.e., harder-to-crack passwords? Do these have greater complexity in their molecular mechanisms? Another open question is do pathogens launch attacks similar to those seen in computer science, e.g., denial-of-service attacks? What are biological equivalents of two-factor authentication?

While I have not presented evidence for cell passwords, encryption, or other security measures, I suggest that they may exist and provide fragments of theory and criteria that the community can use to look for patterns that may demonstrate their existence. There are several examples showing how to quantify Shannon entropy in biochemical systems and protein complexes, reviewed by Waltermann and Klipp^6^. I also have not addressed the critical question of noise in biological systems and measurements, which add considerable complexity to information theoretic analysis of biological systems, which confounds the criterion of whether encryption is present in a signaling cascade because both increase the entropy. If there is encryption and decryption, identifying the molecular mechanisms by which it occurs could yield new and powerful insights into signaling, pathogens, and pathologies.

## Declarations

### Author contributions

A.R. wrote the paper.

## Acknowledgments

The author thanks H. Alexander Ebhardt for critical comments and discussion. The author also thanks two anonymous reviewers for critical comments and suggestions, including bringing to his attention the encrypted genomes of *Oxytricha* and *Stylonychia*.

## Consent for publication

No humans were actively recruited to the work presented here.

## Ethics approval and consent to participate

No humans were actively recruited to the work presented here.

## Publisher’s Note

Springer remains neutral with regard to jurisdictional claims in published maps and institutional affiliations.

